# Contribution of protein conformational heterogeneity to NMR lineshapes at cryogenic temperatures

**DOI:** 10.1101/2023.01.24.525358

**Authors:** Xu Yi, Keith J. Fritzsching, Rivkah Rogawski, Yunyao Xu, Ann E. McDermott

**Author notes:** **Author Contributions:** X.Y., and A.E.M. designed research; X.Y. and K.J.F. performed research; X.Y. and A.E.M. analyzed data; and X.Y. and A.E.M. wrote the paper.

## Abstract

While low temperature NMR holds great promise for the analysis of unstable samples and for sensitizing NMR detection, spectral broadening in frozen protein samples is a common experimental challenge. One hypothesis explaining the additional linewidth is that a variety of conformations are in rapid equilibrium at room temperature and become frozen, creating an inhomogeneous distribution at cryogenic temperatures. Here we investigate conformational heterogeneity by measuring the backbone torsion angle (Ψ) in *E. coli* DHFR at 105K. Motivated by the particularly broad N chemical shift distribution in this and other examples, we modified an established NCCN Ψ experiment to correlate the chemical shift of N_i+1_ to Ψ_i_. With selective ^15^N and ^13^C enrichment of Ile, only the unique I60-I61 pair was expected to be detected in ^13^C’-^15^N correlation spectrum. For this unique amide we detected three different conformation basins based on dispersed chemical shifts. Backbone torsion angles Ψ were determined for each basin 114 ± 7 for the major peak, and 150 ± 8 and 164 ± 16° for the minor peak as contrasted with 118 for the X-ray crystal structure (and 118-130 for various previously reported structures). These studies support the hypothesis that inhomogeneous distributions of protein backbone torsion angles contribute to the lineshape broadening in low temperature NMR spectra.

**Significance Statement:** Understanding protein conformational flexibility is essential for insights into the molecular basis of protein function and the thermodynamics of proteins. Here we investigate the ensemble of protein backbone conformations in a frozen protein freezing, which is likely a close representation for the ensemble in rapid equilibrium at room temperature. Various conformers are spectrally resolved due to the exquisite sensitivity of NMR shifts to local conformations, and NMR methods allow us to directly probe the torsion angles corresponding to each band of chemical shifts.

## Introduction

Many biophysical techniques, such as X-ray crystallography (1) and cryo-EM (2), utilize low temperatures to reduce thermal motion and reduce irradiation damage duering structure determination. Accordingly, the use of low temperatures has allowed investigations of systems that were previously unstable or inaccessible, or improved the precision of structural studies. Decreasing the temperature of samples has also been shown to be beneficial in NMR experiments. At cryogenic temperatures, larger polarizations and improved detection sensitivities can be obtained. Low temperature also facilitates transfer of polarization from electron spins to nuclear spins (3) in a technique known as dynamic nuclear polarization (DNP) which can theoretically enhance the signal to noise ratio up to 660 fold.

In some cases, the use of low sample temperatures for NMR is often accompanied by significant line broadening. Several hypotheses have been advanced to explain this lineshape. A correlation between solvent exposure and broad lineshape suggests a role for solvent interactions in the line broadening; specifically the mean ^15^N linewidths in the DNP spectra of the Pf1 bacteriophage were found to be significantly different between hydrated (mean of 8 ppm, standard deviation of 1.4 ppm) vs non-hydrated residues (mean of 6 ppm and standard deviation of 1.7 ppm) (4). Alternatively, low-temperature broadening can result from exchange dynamics (5). Line broadening might result from the effects of the radicals used for DNP (6, 7), or from difficulties in optimizing cryogenic experimental protocols (8) including the use of relatively low spinning frequencies (4, 9). Several studies indicate that linewidths at low temperatures are typically primarily inhomogeneous. Many studies implicate conformational heterogeneity as an important source of line broadening in low temperature NMR (10–14). In this hypothesis, rapid variation in various structural parameters occurs at room temperature due to conformational fluctuations. Therefore, experimental chemical shifts and other properties are weighted averages of the ensemble. By contrast, freezing the sample results in a static inhomogeneous distribution of conformations and of chemical shifts. We explore this hypothesis to identify some of the conformational degrees of freedom that have broad distributions.

Of the many degrees of freedom in the protein conformation landscape, backbone torsion angles (15–17) are of particular interest and can be accessed by solid state NMR experiments. In this study, precise restraints for Ψ (18) are obtained through dephasing the C’-Cα double-quantum coherence with ^15^N dipolar couplings. The experiment involves detecting the 2D DQ-SQ ^13^C-^13^C correlation spectrum and recording the intensities of the crosspeaks as a function of C-N dephasing times. Sensitivity of this dephasing profile to the relative orientation of the C-N vectors allows a precise determination of torsion angles. This experiment, referred to as the “NCCN Ψ” measurement, has been initially demonstrated for crystalline (^13^C,^15^N-Gly)-(^15^N-Gly)-Gly·HCl (19) and subsequently demonstrated on or applied to other peptides, such as MLF (20) and GNNQQNY fibrils from the yeast prion protein Sup35p (21), as well as SH3 protein residues (18) and β-hairpins in huntingtin exon1 fibrils (22). Here we probe a backbone torsion angle Ψ at low temperature using a modified version of the “NCCN Ψ” measurement (17, 19) to correlate the dipolar angle restraint to the ^15^N shift.

*E. coli* DHFR is a globular enzyme that is very well-studied from the point of view of conformational dynamics and is an important antibiotic target. DHFR catalyzes the reduction of dihydrofolate (DHF) to tetrahydrofolate (THF) using NADPH as a cofactor. There are a number of intermediates in the catalytic cycle (**Fig. 1a**), including: DHFR:NADPH, DHFR:NADPH:DHF, DHFR:NADP+:THF, DHFR:THF and DHFR:NADPH:THF. To accommodate these chemical changes, DHFR accesses a variety of conformations, including two referred to as closed and occluded (**Fig. 1b**). Trimethoprim (TMP) is an effective antibiotic and has a very high binding affinity (K_d_ = 6 pM) (23) to *E. coli* DHFR competing for the substrate and appears to trap the protein in a closed conformation (**Fig. 1c**). The protein contains a cofactor binding pocket and a substrate binding pocket which both undergo intermediate exchange behavior at room temperature. A unique Ile-Ile amino acid pair (60-61) located in a beta-sheet structure close to the cofactor binding pocket is the subject of this study. The value of Ψ for residue 60 is 118°according to the X-ray crystallography structure of *E. coli* DHFR:TMP (24) and it varies from 118°to 130°among the structures of its native complexes in the catalytical cycle (25), as listed in **Table 1S**.

**Fig. 1.**
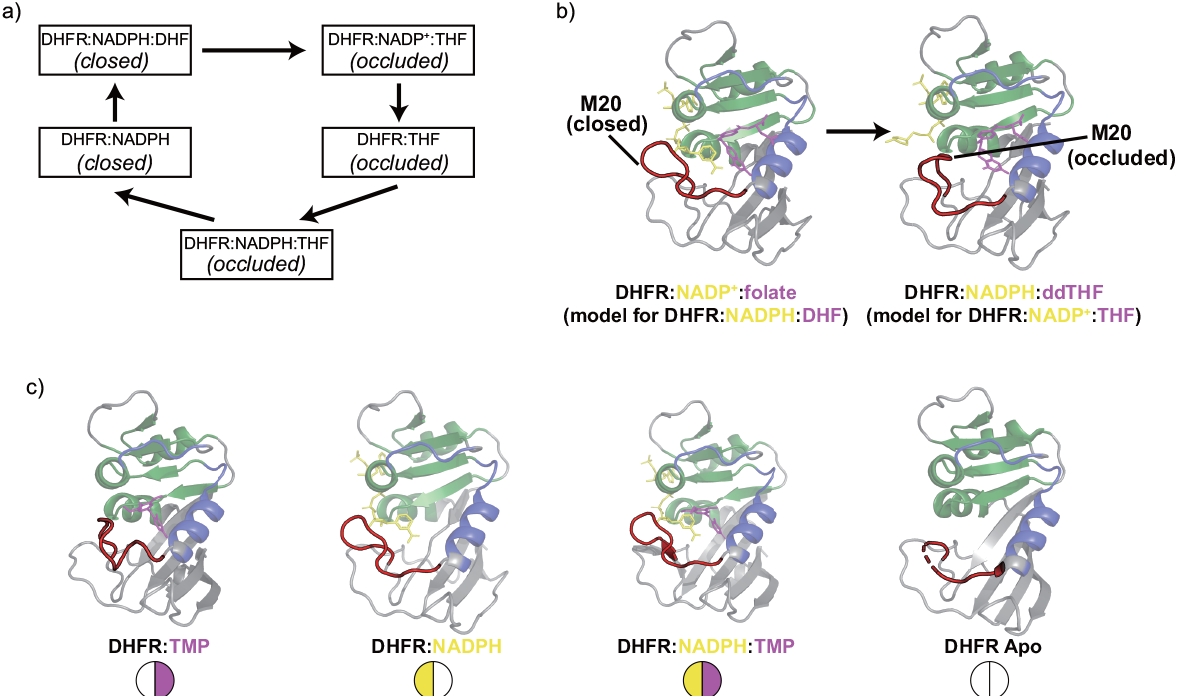
(a) Conformational changes during the DHFR catalytic cycle (25). (a) DHFR catalyzes the reduction of dihydrofolate (DHF) to tetrahydrofolate (THF) in cells. Five key complexes are proposed in the *E. coli* DHFR catalytic cycle. The DHFR:TMP state is considered analogous to DHFR:THF state. (b) The conformation of the M20 loop and other regions change from closed to occluded during hydride transfer. Residues that define the active site loop conformation, substrate-binding, and cofactor-binding markers are colored red, blue, and green, respectively. The substrates are colored magenta and the cofactors are colored yellow. The design of the figures in (a) and (b) comes from previous work (25, 26). (c) The DHFR:TMP (T) (24) state and three associated DHFR liganded states are analyzed using solution-state NMR, including DHFR:NADPH (N) (PDB: 1rx1) state, DHFR:NADPH:TMP (NT) state and the Apo (A) (PDB: 5dfr) state. Because the DHFR:NADPH:TMP (NT) state has no crystal structure available, a model is shown created by docking TMP into the DHFR:NADPH (N) (PDB: 1rx1) structure.

## Results

### Selective isotopic enrichment scheme

To explore the effects of conformational heterogeneity on lineshape, in view of the relatively broad lines, we designed experiments involving selective isotopic enrichment that would highlight a mixture of putatively static and dynamic sites in ^13^C’-^15^N correlation spectra. To that end, we prepared ^13^C,^15^N-Ile, ^15^N-Gly labeled *E. coli* DHFR:TMP (IG-DHFR), for which 5 sites will have ^13^C’-^15^N directly bond pairs detected in ^13^C,^15^N correlation spectrum, specifically a selective DCP experiment with short contact time based on *E. coli* DHFR sequence: Ile2-Ser3 (sheet), Ile14-Gly15 (coil), Ile50-Gly51 (helix), Ile60-Ile61 (sheet), Ile94-Gly95 (coil), shown in **Fig. 2a**. Several regions in *E. coli* DHFR are expected to be mobile in the mechanism, and are in fact notably flexible based on comparing torsion angles in different DHFR ligand-bound structures. For example, sites in the M20 loop (especially Ile14 and Pro21) and the GH loop (including Ser148) vary significantly in their conformations for different complexes (24, 25) (see **Table 1S)**. Regarding residues observed in our experiment, the torsion angle variation of Ile14-Gly15 in the mobile M20 loop is up to 187°; Ile14 is believed to flip (I14Ψ = −11°in DHFR:NADPH:DHF; I14Ψ = 157°in DHFR:NADP+:THF) during the transition from the closed to occluded conformations (25). In the DHFR:TMP complex, both Ile50 and Ile94 are close to the TMP ligand and are possibly affected by the flipping of the TMP trimethoxybenzyl ring. By contrast, Ile2-Ser3 and Ile60-Ile61 are expected to be immobile based on X-ray crystallography. Moreover, in support of this prediction, solution-state NMR T2 relaxation measurements of DHFR complexes and studies of its catalytical cycle, Gly15, Gly51 and Gly95 were observed to exhibit intermediate exchange behavior at room temperature (for the amidic NH group), while Ser3 and Ile61 were not (26). Using this isotope enrichment scheme, we explored whether there is any relation between the low-temperature linewidths and the previous indications for conformational dynamics.

**Fig. 2.**
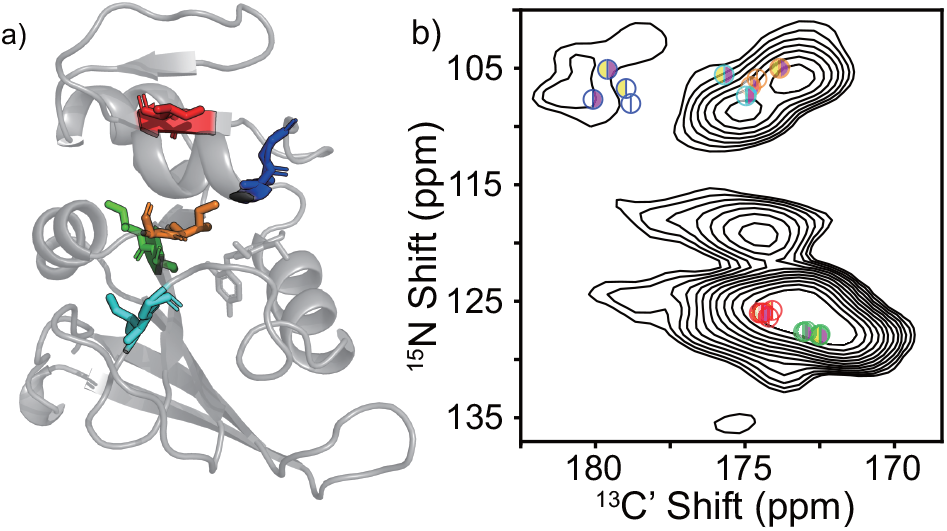
(a) Cartoon of the structure of *E. coli* DHFR:TMP (24) with pairs detected in low temperature NMR spectra of the IG-DHFR highlighted with stick rendering of the sidechains: I2-S3 (green), I14-G15 (cyan), I50-G51 (blue), I94-G95 (orange) and I60-I61 (red). (b) N-C’ correlated DCP spectra of IG-DHFR with likely assignments indicated. The details of spectra collection and processing are listed in **Table 2S**. The chemical shift assignments for 4 liganded states of *E. coli* DHFR are shown with different markers as **Fig. 1c** and colored accordingly as (a). Significant overlap between these peaks introduces some ambiguity in these assignments as well as uncertainty in determining the lineshape of each individual site. Therefore, we next studied a sample where a single amide bond could be detected by low temperature NMR.

In a separate experiment, we prepared ^13^C,^15^N-Ile labeled *E. coli* DHFR:TMP (I-DHFR), in which the only ^13^C’-^15^N directly bonded pair expected to be detected in a selective ^15^N-^13^C correlation experiment is the amide connecting the only sequential I-I pair, residues 60-61. With this sample, we explored in detail the flexibility of a single site and the relation to low-temperature linewidths, without concern about spectral overall with other sites.

### Conformational homogeneity in solution-state NMR spectra

To prepare homogeneous *E. coli* DHFR samples in the desired ligand-bound state we removed residual endogenous THF by an unfolding-refolding procedure after purification. Solution NMR spectra of refolded ^13^C,^15^N-Ile labeled *E. coli* DHFR:TMP (I-DHFR) (**Fig. S2)** were compared to spectra from uniformly ^13^C,^15^N enriched samples from which assignments of 4 different liganded states were determined including DHFR:TMP (T), DHFR:NADPH (N), DHFR:NADPH:TMP (NT) and Apo (A) state. All 12 Ile in the *E. coli* DHFR sequence were observed in the solution ^15^N-HSQC spectra of I-DHFR. Chemical shifts and signal intensities match expectations based on with the spectra of uniformly ^13^C,^15^N T state (**Fig. S2a**), confirming successful selective enrichment and refolding. The same protein sample was used for low-temperature DNP-enhanced SSNMR spectroscopy (**Fig. 3** and **S5)**. Though the ligation of this I-DHFR sample is homogeneous (namely saturated with respect to TMP ligand and with no cofactor or other ligand bound) some Ile residues exhibit minor peaks suggesting a mixture of conformers. Ile61 (**Fig. S2b-d**), Ile155 (**Fig. S2a**) and Ile41 (**Fig. S2a**) have minor peaks with similar chemical shifts as either DHFR:NADPH:TMP (NT) state or DHFR:NADPH (N) state. Possibly the minor peaks result from slow chemical exchange to other conformations.

**Fig. 3.**
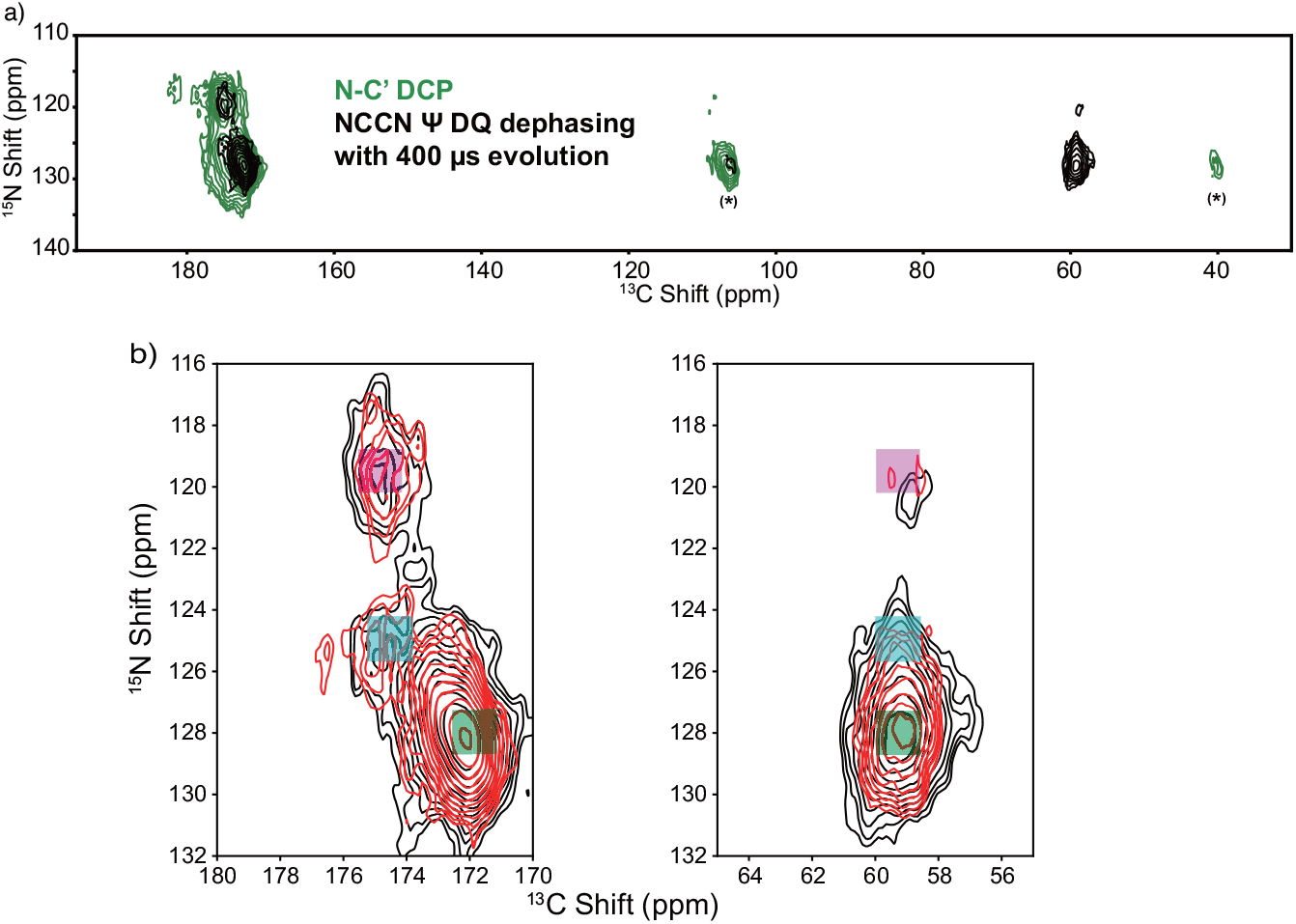
DNP spectra of ^13^C,^15^N-Ile DHFR:TMP (I-DHFR) collected at 105 K. (a) The N-C’ correlation spectrum of I-DHFR (green) collected with DCP pulse sequence (37, 38) is compared with the S_0_ spectrum with 400 μs evolution (t_CN_ = 4 τr) of the NCCN Ψ DQ dephasing experiment (black). No Cα intensity is observed after the DCP transfer (green), which is spectrally selective from Ni+1 to Ci. The asterisks (*) are the first and the second spinning sidebands of the C’. In the NCCN Ψ DQ dephasing experiment, the ^13^Cαi-^13^C’i DQ coherence was reconverted to both Cα and C’ with SPC5 (33), resulting in the observed Cα peaks (black). (b) The zoomed-in S_0_ spectra (black) and S_r_ spectra (red) with 400 μs evolution at NC’ and NCα regions. Three peaks at N chemical shifts of 128 ppm, 125 ppm and 119.5 ppm are highlighted with green (Peak 1), cyan (Peak 2) and purple (Peak 3) square overlaid ornaments. The details of spectral collection and processing are listed in **Table 2S**.

### Lineshape broadening in DNP-enhanced NMR spectra of frozen solutions of DHFR

5 amide pairs are expected to appear in DNP-enhanced N-C correlation spectra of IG-DHFR, and 5 major peaks are indicated on the experimental data (**Fig. 2b**). They are labeled based comparison to solution-state NMR chemical shifts, though peak overlap and minor additional peaks cause these tentative assignments. The ^15^N amide linewidth of these residues varies from 3.4 ppm to 8.2 ppm (**Fig. S3**), while the ^13^C’ linewidth varies from 1.9 ppm to 2.3 ppm. There is no apparent relationship between the magnitude of linewidth at low temperature and indications of chemical shift variation due to conformational dynamics at room temperature from solution-state NMR studies (**Fig. S4a**). Several ^15^N linewidths are considerably larger than the range in chemical shift expected for the various species in exchange in solution (26) (less than 2 ppm), and the two quantities are apparently uncorrelated. Thus, the variation in isotropic shift observed in DNP is probably not specifically due to the millisecond and microsecond conformational exchange processes observed in solution NMR in this example.

### Torsion angle measurements

In order to explore the conformational heterogeneity within a single confidently assigned amino acid residue, we characterized refolded ^13^C, ^15^N-Ile labeled *E. coli* DHFR:TMP (I-DHFR) using chemical shift and torsion angle measurements, probing the unique II pair in DHFR (60-61). To test whether the broad spectra lines in the low temperature spectra of DHFR reflect an inhomogeneous mixture of various conformers, restraints on the torsion angle Ψ were obtained using the NCCN Ψ DQ dephasing experiment (18, 19) (illustrated in **Fig. S1**), modified to be a pseudo 3D experiment, with chemical shifts for N and C’ measured in a 2D spectrum, and dephasing for torsion angle determination in the third dimension.

We investigated whether distinct chemical shifts in the broad low temperature peak are associated with distinct conformations by comparing the dephasing profiles at various positions in the N-C correlation spectrum. The 2D N-C’ correlation spectrum is shown in **Fig. 3a** (green). The chemical shift range of I61N is 115 ppm - 135 ppm and the I60C’ chemical shift range is 170 ppm - 180 ppm. The extremely broad peak of I60C-I61N appears to be organized in multiple “basins”, suggesting the existence of various conformational basins (collections of similar structures). The N-C correlation spectrum with the shortest evolution time is shown in **Fig. 3** (black). The control experiment, S_0_ without dephasing (**Fig. 3a**, black) is compared to the simple DCP 2D spectrum (**Fig. 3a**, green) and has a similar peak shape with approximately 15% of the signal intensity, since losses occur when exciting DQ coherences and then reconverting back to SQ coherence. Considering the peak shape of I60C-I61N, three peak regions were defined for torsion angle analysis (**Fig. 3b**), Peak 1 (green), Peak 2 (cyan), Peak 3 (purple). The normalized intensity ratios of each dephased spectrum (pulse sequence **Fig. S1c**) and its corresponding reference spectrum (pulse sequence **Fig. S1d**) (S_r_/S_0_ defined in Eq. 2 in SI) were extracted from all the 2D spectra with evolution times varying from 4 *τr* to 14 *τr* (**Fig. S5**). The intensities are plotted as a dephasing curve for torsion angle analysis (**Fig. 4**), with each of the three “subpeaks” of I60 of DHFR:TMP indicated separately (dephasing times as listed in the figure caption). The data from the NCCN Ψ DQ dephasing experiment indicate that at least three subpeaks in the I60 spectrum have different dephasing time profiles. Using a Chi-square test to fit the three peaks, there appear to be three conformations present with Ψ =114 ± 7, 150 ± 8 and 164 ± 16°, respectively. The error bar of the Chi-square fit is determined by a 90% confidence level. Details of the NCCN Ψ DQ dephasing pulse sequences and curve fittings are included in SI.

**Fig. 4.**
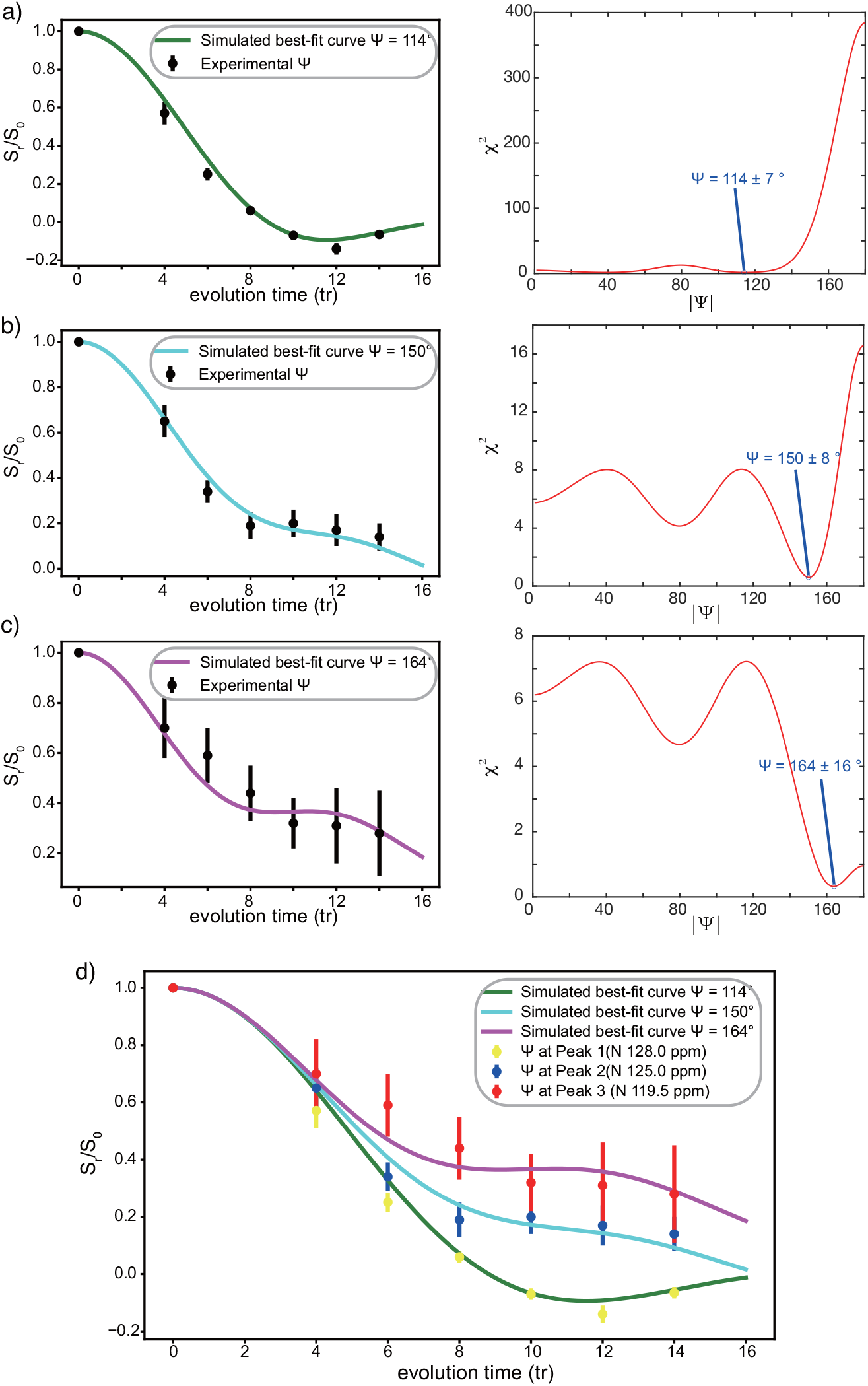
Dipolar dephasing of (a) Peak 1 (N 128.0 ppm, green in **Fig. 3b**), (b) Peak 2 (N 125.0 ppm, cyan in **Fig. 3b**) and (c) Peak (N 119.5 ppm, magenta in **Fig. 3b**) with corresponding best-fit curves based on a Chi-square test. The normalized intensity ratio S_r_/S_0_ was calculated from peak intensities in S_0_ and S_r_ spectra (**Fig. S5**) collected using the NCCN Ψ DQ dephasing experiment (18, 19) (illustrated in **Fig. S1**). The overlap of the three decay curves is displayed in (d), which clearly shows the different decay time profiles for the three peaks.

The X-ray crystallography of *E. coli* DHFR: TMP shows that the Ile60 and Ile61 are located in a beta-sheet with Ψ for I60 of 118°, which is within the error bar range of Peak 1. The X-ray structures of *E. coli* DHFR complexed in various liganded states show I60Ψ with a variation range from 118°to 130°. The two complexes with the closed conformation in *E. coli* DHFR catalytic cycle exhibit a Ψ angle of 120°and 122°at Ile60. Among structures of the native protein complexes the value of Ψ for residue 60 varies up to 130°.

Despite the fact that the range of torsion angle values indicated by these SSNMR data exceed that indicated in the crystal structures, it is noteworthy that the values indicated by the SSNMR data are all compatible with the idea that the residue remains in the beta basin. The range of chemical shifts also indicate beta structures. The three peaks observed at low temperature have C’ and Ca chemical shifts compatible with typical sheet or random coil values. The average N chemical shift for Ile in beta-sheet conformations is 124 ± 5 ppm, while that for a random coil is 121 ± 5 ppm (27), indicating that the two stronger peaks (Peak 1 and Peak 2 with N chemical shift of 125 ppm and 128 ppm) are likely to be a beta-sheet structure. Peak 1 has a N shift of 119.5 ppm, which is more likely to be a random coil but compatible with sheet.

The average properties indicated by crystallography or solution NMR agree well with the average of the three peaks seen at low temperature. Weighted averages of the torsion angle values determined for the three low temperature SSNMR peaks is in the range of the values from crystallography (128°) and a typical value for beta sheets. The average of the chemical shifts in the low temperature SSNMR data, using weighted sum of the three peaks were also in good agreement with the solution NMR shifts. Table S4 analyses three shifts in this fashion: Ile61N, Ile60C’ and Ile60Cα. Overall, it is reasonable to hypothesize that the solution ensemble includes a rapid average of three states corresponding to the three peaks observed at low temperature.

We eliminated the possibility of irreversible sample alteration during DNP sample preparation and freezing. After the low-temperature experiments, the DNP sample was collected and buffer exchanged and characterized with room temperature solution-state NMR experiments. Comparing the ^15^N-HSQC spectra before and after DNP measurements (**Fig. S6**), all 12 Ile residues were observed and preserved at the same peak positions and approximate relative intensities. The sample concentration calculation based on the spectral intensity indicates that 85% of the protein sample was successfully recovered for this analysis, and we conclude that the structures we observed at the low temperature result from reproducible conformational basins reversibly trapped by freezing, rather than degradation of the protein.

Since Peak 1 (with ^15^N chemical shift of 128 ppm) is intense and broad, we asked if there exist multiple conformations within this broad peak, each with its own torsion angle. The analysis is shown in **Fig. S7**. Three peak regions were picked based on the peak shape of the Sr spectra after 400 μs (**Fig. S7**): Peak 1A (N 127 ppm), Peak 1B (N 129 ppm), Peak 1C (N 130 ppm). Applying the Chi-square fit of these three peaks, Peak 1A gives Ψ =115 ± 8°; as for Peak 1B and Peak 1C, the Ψ angle corresponding to the minimum Chi-square is 116 and 115°, but in both cases, the minimum chi squared value exceed the range for the 90% confidence interval (Chi-square of Peak 1B is 5.3 and Peak 1C is 3.3), indicating that neither Peak 1B nor Peak 1C is compatible with a different torsion angle value. This result indicates that one torsion angle Ψ well explains this broad peak, and the line broadening within peak 1 may be associated with other structural or spectroscopic parameters.

## Discussion

The torsion angle variation observed for residue 60 at cryogenic temperatures has a range of values that is larger than the variation values reported in various X-ray structures including various ligated states. Such a variation was not anticipated based on the thermal disorder in the structures, since the X-ray crystallography structures of either closed (PDB: 1rx2) or occluded (PDB: 1rx6) conformations of *E. coli* DHFR report B-factors smaller than 15 Å^2^ for backbone atoms in I60-I61 (25) suggesting good precision and rigidity. From the point of view of millisecond and microsecond dynamics and line broadening in solution NMR at room temperature, I60 is not expected to be an exceptionally flexible residue (26, 28), without pronounced relaxation dispersion observed in native ligated states; by all indications, it is a structured residue within a beta-strand (24–26). Notably, low temperature linewidths for this residue are comparable to those from many other residues (7) and other samples (4, 29, 30). Therefore, we hypothesize that structural variations such as large backbone torsion angles likely occur for numerous other backbone sites, while the protein remains stably folded. One possible interpretation is that the variation in (or interconversion between) Peaks 1,2 and 3 also involves correlated motions of other degrees of freedom within the protein, for example, ɸ of the succeeding residue, so that overall the backbone shape is not significantly displaced. In this case, a “rocking” motion of the amide plane within a stably folded protein would occur. Similar motions of the backbone have been discussed previously based on NMR and dynamics simulations (31, 32) and in terms of their significance for protein structural determination (33, 34).

The timescale of interconversion between structures associated with Peak 1, Peak 2 or Peak 3 is not known directly from these data. The narrow solution NMR linewidths for this residue (and others) observed at room temperature suggest that these structures (corresponding to Peaks 1, 2 and 3) interconvert on a sub nanosecond timescale. Fast timescale backbone and sidechain dynamics of DHFR (~50 ps to 10 ns) have been studied using solution NMR indicating S^2^ > 0.8 for the NH vector (36). This relatively rigid description was determined to be consistent with the diffraction data for the crystalline protein (35). The present cryogenic SSNMR data suggest a somewhat lower order parameter for Ile60 (considering a two site model with 30% of the population at an angle of 40°, giving S^2^ ~0.65). This suggests that these SSNMR data may reflect dynamics on an even faster timescale than detected by solution NMR (low ps), consistent with some previous studies (31, 32). By contrast, the inhomogeneously broad lines in cryogenic NMR spectra suggest that the structures interconvert on a timescale of many ms or slower at 105 K. If these conjectures are correct, a very steep change in the rate constant with respect to the temperature suggests that the solvent (and its phase change) plays an important role.

In conclusion, we determined the range of backbone torsion angle Ψ for a specific residue in a frozen protein, DHFR, at 105K using a modified NCCN solid-state NMR dihedral angle measurement. We showed that the broad NMR peak for the single amide N-C’ correlation of I60-I61 contains multiple conformational subspecies with distinct chemical shifts and torsion angles. This result supports the hypothesis that the origin of broad lines in NMR of frozen samples is dominated by conformational heterogeneity, including backbone torsion angle heterogeneity, at least for some protein sites.

## Acknowledgments

This work was supported by a grant from the NSF (MCB 1913885), and a Biomedical Technology Development and Dissemination Center grant from the NIH (1RM1GM145397-01). We thank Dr. Michael J. Goger, Dr. Eric Keeler and Dr. Shibani Bhattacharya of the New York Structural Biology Center (NYSBC) for help with instrumentation. We thank Kirk Baughman and Chengming He for their advice for both experiments and manuscripts.

## Notes

The authors declare no conflict of interest.

**Competing Interest Statement:** Disclose any competing interests here.

### Competing Interest Statement

The authors have declared no competing interest.

## References

1. A. E. Goeta, J. A. K. Howard, Low temperature single crystal X-ray diffraction: Advantages, instrumentation and applications. Chem. Soc. Rev. 33, 490–500 (2004).

2. X. Bai, G. McMullan, S. H. W. Scheres, How cryo-EM is revolutionizing structural biology. Trends Biochem. Sci. 40, 49–57 (2015).

3. P. Wenk, M. Kaushik, D. Richter, Dynamic nuclear polarization of nucleic acid with endogenously bound manganese. 97–109 (2015).

4. I. V. Sergeyev, B. Itin, R. Rogawski, L. A. Day, A. E. McDermott, Efficient assignment and NMR analysis of an intact virus using sequential side-chain correlations and DNP sensitization. Proc. Natl. Acad. Sci. U. S. A. 114, 5171–5176 (2017).

5. V. S. Bajaj, P. C. A. A. van der Wel, R. G. Griffin, Observation of a Low-Temperature, Dynamically Driven Structural Transition in a Polypeptide by Solid-State NMR Spectroscopy. J. Am. Chem. Soc. 131, 118–128 (2009).

6. I. V. Sergeyev, C. M. Quinn, J. Struppe, A. M. Gronenborn, T. Polenova, Competing transfer pathways in direct and indirect dynamic nuclear polarization magic anglespinning nuclear magnetic resonance experiments on HIV-1 capsid assemblies: implications for sensitivity and resolution. Magn. Reson. 2, 239–249 (2021).

7. R. Rogawski, et al., NMR Signal Quenching from Bound Biradical Affinity Reagents in DNP Samples. J. Phys. Chem. B 121, 10770–10781 (2017).

8. K. J. Fritzsching, B. Itin, A. E. McDermott, N,N-Diethylmethylamine as lineshape standard for NMR above 130 K. J. Magn. Reson. 287, 110–112 (2018).

9. K. Jaudzems, et al., Dynamic Nuclear Polarization-Enhanced Biomolecular NMR Spectroscopy at High Magnetic Field with Fast Magic-Angle Spinning. Angew. Chemie - Int. Ed. 57, 7458–7462 (2018).

10. Y. Su, M. Hong, Conformational disorder of membrane peptides investigated from solidstate NMR line widths and line shapes. J. Phys. Chem. B 115, 10758–10767 (2011).

11. R. Gupta, et al., Dynamic Nuclear Polarization Magic-Angle Spinning Nuclear Magnetic Resonance Combined with Molecular Dynamics Simulations Permits Detection of Order and Disorder in Viral Assemblies. J. Phys. Chem. B 123, 5048–5058 (2019).

12. M. J. Bayro, et al., Intermolecular structure determination of amyloid fibrils with magic-angle spinning and dynamic nuclear polarization NMR. J. Am. Chem. Soc. 133, 13967–13974 (2011).

13. L. Reggie, J. J. Lopez, I. Collinson, C. Glaubitz, M. Lorch, Dynamic nuclear polarization-enhanced solid-state NMR of a 13C-labeled signal peptide bound to lipid-reconstituted sec translocon. J. Am. Chem. Soc. 133, 19084–19086 (2011).

14. J. M. Lopez Del Amo, D. Schneider, A. Loquet, A. Lange, B. Reif, Cryogenic solid state NMR studies of fibrils of the Alzheimer’s disease amyloid-β peptide: Perspectives for DNP. J. Biomol. NMR 56, 359–363 (2013).

15. X. Feng, et al., Direct determination of a molecular torsional angle by solid-state NMR. Chem. Phys. Lett. 257, 314–320 (1996).

16. P. R. Costa, J. D. Gross, M. Hong, R. G. Griffin, Solid-state NMR measurement of Ψ in peptides: a NCCN 2Q-heteronuclear local field experiment. Chem. Phys. Lett. 280, 95–103 (1997).

17. V. Ladizhansky, C. P. Jaroniec, A. Diehl, H. Oschkinat, R. G. Griffin, Measurement of Multiple ψ Torsion Angles in Uniformly 13 C, 15 N-Labeled α-Spectrin SH3 Domain Using 3D 15 N- 13 C- 13 C- 15 N MAS Dipolar-Chemical Shift Correlation Spectroscopy. J. Am. Chem. Soc. 125, 6827–6833 (2003).

18. V. Ladizhansky, C. P. Jaroniec, A. Diehl, H. Oschkinat, R. G. Griffin, Measurement of multiple ψ torsion angles in uniformly 13C,15N-labeled α-spectrin SH3 domain using 3D 15N-13C--13C-15N MAS dipolar-chemical shift correlation spectroscopy. J. Am. Chem. Soc. 125, 6827–6833 (2003).

19. P. R. Costa, J. D. Gross, M. Hong, R. G. Griffin, Solid-state NMR measurement of Ψ in peptides: A NCCN 2Q-heteronuclear local field experiment. Chem. Phys. Lett. 280, 95–103 (1997).

20. C. M. Rienstra, et al., De novo determination of peptide structure with solid-state magic-angle spinning NMR spectroscopy. Proc. Natl. Acad. Sci. 99, 10260–10265 (2002).

21. P. C. A. Van Der Wel, J. R. Lewandowski, R. G. Griffin, Structural characterization of GNNQQNY amyloid fibrils by magic angle spinning NMR. Biochemistry 49, 9457–9469 (2010).

22. C. L. Hoop, et al., Huntingtin exon 1 fibrils feature an interdigitated β-hairpin-based polyglutamine core. Proc. Natl. Acad. Sci. U. S. A. 113, 1546–1551 (2016).

23. S. P. Sasso, R. M. Gilli, J. C. Sari, O. S. Rimet, C. M. Briand, Thermodynamic study of dihydrofolate reductase inhibitor selectivity. Biochim. Biophys. Acta (BBA)/Protein Struct. Mol. 1207, 74–79 (1994).

24. R. Rogawski, Dynamic Nuclear Polarization with Biradical Affinity Reagents (2018).

25. M. R. Sawaya, J. Kraut, Loop and Subdomain Movements in the Mechanism of Escherichia coli Dihydrofolate Reductase: Crystallographic Evidence,. Biochemistry 36, 586–603 (1997).

26. D. D. Boehr, D. McElheny, H. J. Dyson, P. E. Wright, The Dynamic Energy Landscape of Dihydrofolate Reductase Catalysis. Science (80-.). 313, 1638–1642 (2006).

27. Y. Wang, Probability-based protein secondary structure identification using combined NMR chemical-shift data. Protein Sci. 11, 852–861 (2002).

28. D. D. Boehr, D. McElheny, H. J. Dyson, P. E. Wright, Millisecond timescale fluctuations in dihydrofolate reductase are exquisitely sensitive to the bound ligands. Proc. Natl. Acad. Sci. 107, 1373–1378 (2010).

29. A. Potapov, W. M. Yau, R. Ghirlando, K. R. Thurber, R. Tycko, Successive Stages of Amyloid-β Self-Assembly Characterized by Solid-State Nuclear Magnetic Resonance with Dynamic Nuclear Polarization. J. Am. Chem. Soc. 137, 8294–8307 (2015).

30. S. Y. Liao, M. Lee, T. Wang, I. V. Sergeyev, M. Hong, Efficient DNP NMR of membrane proteins: Sample preparation protocols, sensitivity, and radical location. J. Biomol. NMR 64, 223–237 (2016).

31. J. C. Williams, A. E. McDermott, Variable NMR spin-lattice relaxation times in secondary amides: Effect of ramachandran angles on librational dynamics. J. Phys. Chem. B 102, 6248–6259 (1998).

32. J. E. Fitzgerald, A. K. Jha, T. R. Sosnick, K. F. Freed, Polypeptide Motions Are Dominated by Peptide Group Oscillations Resulting from Dihedral Angle Correlations between Nearest Neighbors. Biochemistry 46, 669–682 (2007).

33. L. Salmon, et al., Multi-Timescale Conformational Dynamics of the SH3 Domain of CD2-Associated Protein using NMR Spectroscopy and Accelerated Molecular Dynamics. Angew. Chemie Int. Ed. 51, 6103–6106 (2012).

34. D. A. Keedy, et al., Crystal cryocooling distorts conformational heterogeneity in a model michaelis complex of DHFR. Structure 22, 899–910 (2014).

35. R. Bryn Fenwick, H. Van Den Bedem, J. S. Fraser, P. E. Wright, Integrated description of protein dynamics from room-temperature X-ray crystallography and NMR. Proc. Natl. Acad. Sci. U. S. A. 111 (2014).

36. J. R. Schnell, H. J. Dyson, P. E. Wright, Correction to Effect of Cofactor Binding and Loop Conformation on Side Chain Methyl Dynamics in Dihydrofolate Reductase. Biochemistry 52, 2383–2383 (2013).

37. M. Baldus, A. T. Petkova, J. Herzfeld, R. G. Griffin, Cross polarization in the tilted frame: Assignment and spectral simplification in heteronuclear spin systems. Mol. Phys. 95, 1197–1207 (1998).

38. J. Schaefer, R.. McKay, E. O. Stejskal, Double-cross-polarization NMR of solids. J. Magn. Reson. 34, 443–447 (1979).

