## supplemental figures for "Contribution of protein conformational heterogeneity to NMR lineshapes at cryogenic temperatures"

PNAS

**This PDF file includes:**

- Figures S1 to S10
- Tables S1 to S4
- SI References

### Supporting Information Text

#### Materials and Methods

**Selective isotopic enrichment of *E. coli* DHFR:** 50 mL LB mixed with 1% (10 mg/mL) glucose was inoculated with a small shard of ice from DHFR glycerol stock overnight. 2x 500 mL LB with 0.5% (5 mg/mL) glucose was inoculated with 5 mL of the overnight preculture and set to shake at 37 degrees, 250 rpm. Cells were collected when $OD_{600}$ = 0.8 and 0.9, then resuspended into 250 mL M9 buffer with 50 mg each of all 20 amino acids, some of which were labeled. The shaking speed was changed to 300 rpm. After 20 min of shaking, cells were induced using 1M IPTG at a 1000-fold dilution. Cells were harvested after 2 h of expression.

**Protein purification and refolding:** *E. coli* DHFR was purified using FPLC with gradient washing (5 mM-25 mM imidazole) and step elution (200 mM imidazole and 500 mM imidazole). Purified *E. coli* DHFR samples were then unfolded in unfolding buffer(8 M urea, 300 mM NaCl, 75 mM Na2HPO4 , 5 mM BME (2-mercaptoethanol), pH 8) and incubated at room temperature for 1 h. Unfolded DHFR was purified using a gravity column with washing buffer (20 mM imidazole, 6 M urea, pH 8.0) and elution buffer (500 mM imidazole, 6 M urea, pH 7.5). Purified unfolded *E. coli* DHFR was diluted into 25-fold refolding buffer (300 mM NaCl, 75 mM $Na_2HPO_4$ , pH 8.0) with 10x TMP at room temperature and gently stirred overnight at 4 degrees. Native purification using FPLC column was applied once more to make sure samples were free of urea. Elution fractions were transferred into HEPES buffer (70 mM HEPES, 150 mM KCl, 0.1 mM EDTA, 5 mM DTT, pH 7.6).

**DNP experiments:**

Sample preparation: *E. coli* DHFR sample was packed into 1.9 mm rotor with a mixed solvent in the ratio 60/30/10 (v/v/v) $D_2O$/glycerol/$H_2O$, a protein concentration of ~4 mM, AMUPOL concentration of ~10 mM, and saturating concentration of TMP is ~10 mM.

DNP experiments: DNP-SSNMR spectroscopy was carried out at the New York Structural Biology Center on a 14.1-T Bruker AVANCE-III 600/89 spectrometer (600-MHz $^1$H field) equipped with a 395-GHz gyrotron, low-temperature MAS cabinet, and a three-channel $^1$H– $^{13}$C– $^{15}$N (HCN) 1.9-mm DNP probe.

**Solution state NMR experiments:** Refolded $^{13}$C, $^{15}$N-Ile DHFR in HEPES buffer (70 mM HEPES, 150 mM KCl, 0.1 mM EDTA, pH 7.6, with with 5% $D_2O$) was transferred into a 4 mm Shigemi tube for NMR measurement. Solution-state HSQC and H(N)C' and H(NC')Cα spectra were collected at 300 K on a Bruker 700 MHz or 800 MHz $^1$H spectrometer in New York Structural Biology Center.

**NCCN Ψ DQ dephasing experiment:**

In DNP spectra, the N dimension has a particularly broad peak shape. We therefore elected to modify the experimental format to correlate the chemical shift of $N_{i+1}$ to $\Psi_i$, in effect testing whether the poor spectral resolution in the amidic N dimension is associated with the protein conformational heterogeneity. We developed a sequence somewhat analogous to previously reported NC'CA NCCN measurements (1). Our sequence begins by generating $^{15}N_{i+1}$ magnetization (using an H-N CP transfer (2)) and measuring the nitrogen shift. A DCP (3) transfer then moves the magnetization to C' of residue i, and acts as a filter to only observe specific peaks with directly bound $^{15}$N-$^{13}$C pairs. C-C double quantum coherence is then created from SQ C' magnetization using SPC5 (4). Subsequently, the DQ coherence evolves under the influence of both N-C dipolar couplings which are reintroduced using REDOR to probe NCCN torsion angle constraints for residue i (5). The DQ coherence is then converted to detectible SQ coherence on both C' and the Ca of residue i. The resulting N(i+1)-C'(i) correlation spectrum selectively detects specific amino acid pairs, and provides restraints for Ψ(i) in a third pseudo dimension.

The effective Hamiltonian of the "reintroduced" dipolar coupling is:

$H_{REDOR} = \omega_D^{C'_iN_{i+1}} C'_Z N_Z + \omega_D^{C\alpha_iN_i} C\alpha_Z N_Z$ (Eq. 1)

This Hamiltonian generates dephasing behavior which will result in time dependent signal reduction:

$\frac{S_r}{S_0} = < \cos\left(\omega_D^{C'_iN_{i+1}} t_{CN}\right) \cos\left(\omega_D^{C\alpha_iN_i} t_{CN}\right) > = \cos\left(\overline{\omega_D^{C'_iN_{i+1}}}\, t_{CN}\right) \cos\left(\overline{\omega_D^{C\alpha_iN_i}}\, t_{CN}\right)$ (Eq. 2)

The $\omega_D$ includes the torsion angle information because:

$\overline{\omega_D^{CN}} = 2Im(\omega_D^{CN(-1,m)})$ (Eq. 3)

$\omega_D^{CN(-1,m)} = -\frac{\mu_0}{4\pi}\frac{\gamma_C\gamma_N\hbar}{r_{CN}^3}\sum_{m=-2}^{2} D_{0,m}^{(2)}(\Omega_{PM}^{CN})D_{m,-1}^{(2)}(\Omega_{MR})d_{-1,0}^{(2)}(\theta_M)$ (Eq. 4)

$\theta_M$is magic angle ($arctan\sqrt{2}$),. $D_{0,m}^{(2)}(\Omega_{PM}^{CN})$describes transformation of corresponding CN dipole tensor from its principal axis frame to the molecular frame which can be chosen to be the same as principal axis frame of CC dipole tensor. The Euler angles $\Omega_{PM} = (\alpha_{PM}, \beta_{PM}, \gamma_{PM})$ relate these two frames, in which $\beta_{PM}^{C'_iN_{i+1}} - \beta_{PM}^{C\alpha_iN_i}$ is bond angle, and $\gamma_{PM}^{C'_iN_{i+1}} - \gamma_{PM}^{C\alpha_iN_i}$ is torsion angle, as shown in **Fig.S1b**. Another set of Euler angles $\Omega_{MR}$ are random variables that relate the molecular frame to the rotor frame and are sampled to create the powder average.
Simulations of this pulse sequence were carried out in MATLAB_R2019a using spinach_2.1.4400 (6).

**Curve fitting:**

The Chi-square test (7) was applied to determine the best fit and the error limits for acceptable fits (8).

$\chi^2 = \sum_i(\frac{S_i - E_i}{\sigma_i})^2$ (Eq. 5)

$S_i$ are the data points, in this case, the ratio of $S_r$ and $S_0$ in (Eq. 2); $E_i$ are the model predictions- namely, in this case, the theoretical ratio calculated in the simulations; $\sigma_i$ are the standard deviations, which were determined from spectral noise as:

$\sigma_i = \left(\frac{S_r}{S_0}\right)_i \times \left[\left(\frac{1}{Signal\ to\ Noise}\right)_r + \left(\frac{1}{Signal\ to\ Noise}\right)_0\right]_i$ (Eq. 6)

The value for $\sigma_i$ is indicated on each experimental data point in **Fig.4** and **Fig. S7** using error bars.

In order to test for overfitting, reduced Chi-square tests were used. Because the dihedral angle is the only free parameter in the fitting simulation, the number of degrees of freedom is determined based on data points.

In our case, the final formula is

${\chi_v}^2 = \frac{1}{n-1}\sum_i(\frac{S_i - E_i}{\sigma_i})^2$ (Eq. 7)

Here n is the number of data points used for the reduced Chi-square tests.

The pulse sequences illustrated in **Fig. S1** were tested on 2 different peptides: ($^{13}C$,$^{15}N$-Gly)-($^{15}N$-Gly)-Gly·HCl and U-$^{13}C$, $^{15}N$-formyl-Met-Leu-Phe-OH by measuring the Ψ angle of G1 and Met. The normalized intensities were calculated based on the 1D spectra, and the Chi-square tests report accurate fitting results compared to the X-ray structures (**Fig. S8** and **Fig. S9**). Long-range dipolar and J-interaction and CSA tensors have relatively minor effects and the decay curve is dominantly by the relative angle of the two directly-bonded NC dipolar couplings to the two N flanking the C’ and Cα. This result was reproduced in our simulation (**Fig. S10**) and only one-bond NC dipolar couplings were considered in the protein/peptide fittings.

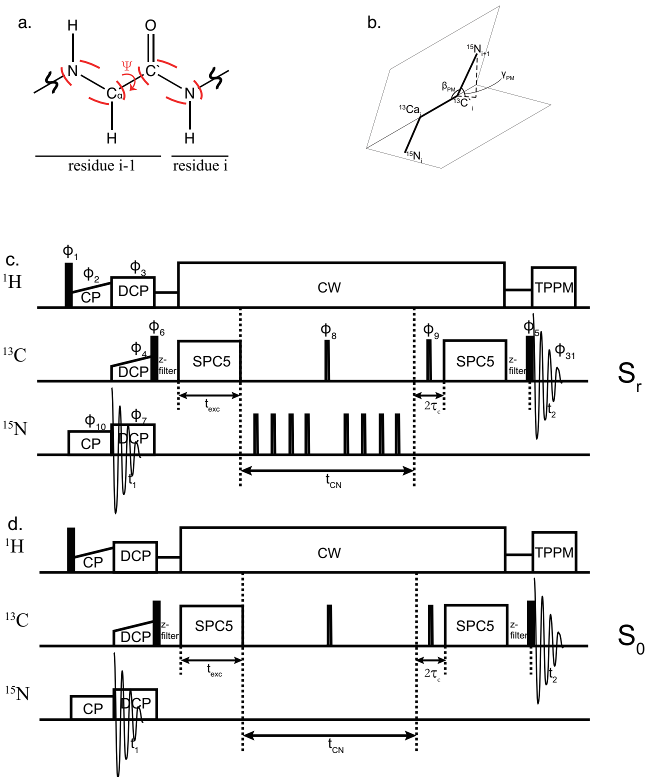


**Fig. S1.** (a) torsion angle psi (Ψ). (b) relationship between Euler angles and torsion angles. (c. d) NCCN Ψ DQ dephasing experiment (1, 9, 10). Closed and open rectangles represent $\pi/2$ and $\pi$ pulses, respectively. Following CP and DCP, DQ coherence is excited during $t_{exc}$ with SPC5 pulse sequence, and then evolve during $t_{CN}$. The DQ coherence is reconverted into SQ before signal acquisition. $t_2$ is the direct dimension detection, $t_1$ is the indirect dimension detection. The phase cycles in c and d were: $\phi_1$= 02, $\phi_2$ = 1, $\phi_3$ = 0, $\phi_4$ = 00002222, $\phi_5$ = 4*{1230} 4*{2301} 4*{3012} 4*{0123}, $\phi_6$ = 31131331, $\phi_7$ = 0, $\phi_8$ = 11221122, $\phi_9$ = 4*{1} 4*{2} 4*{3} 4*{0}, $\phi_{31}$ = 4*{2103} 4*{3210} 4*{0321} 4*{1032}. The $^{15}$N REDOR pulses were phase-cycled according to XY-4 schemes. $\phi_9$ applies the EXOCYCLE to minimize the direct pulsed C signal.

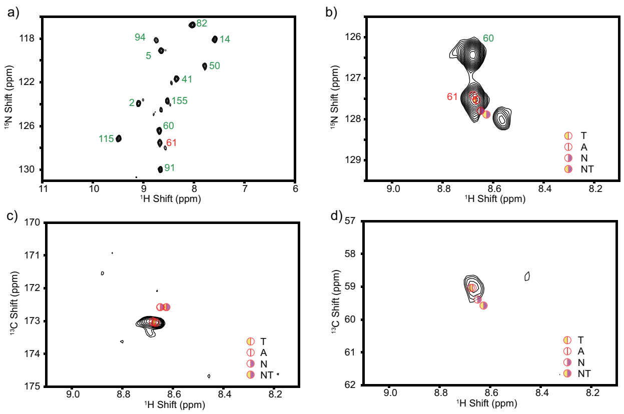

**Fig. S2:** Solution-state NMR spectra of $^{13}C$,$^{15}N$-Ile DHFR:TMP (I-DHFR) collected at 300 K. (a, b) $^{15}N$-HSQC collected with 16 scans per increment in the indirect dimension, and 128 increments in the indirect dimension. All 12 Ile (Ile2N-H, Ile5N-H, Ile14N-H, Ile41N-H, Ile50N-H, Ile60N-H, Ile61N-H, Ile82N-H, Ile91N-H, Ile94N-H, Ile115N-H, Ile155N-H) in *E. coli* DHFR were observed with expected peak intensities compared with uniformly enriched but otherwise similar samples. (c) H(N)C' spectrum and (d) H(NC')Cα spectrum were collected with 256 scans per increment in the indirect dimension, and 64 increments in the indirect dimension. The lowest contour levels in (a) are set to be 5 times of the RMS noise level while 3 times in (b, c, d) and the multiplier is 1.2 for all the spectra. The peaks are in the expected positions based on other 3D backbone correlation and assignment experiments on uniformly enriched but otherwise similar samples. The chemical shift assignments for 4 liganded states of *E. coli* DHFR are shown with different markers as **Fig. 1c**. The acquisition and processing details are listed in **Table 3S**.

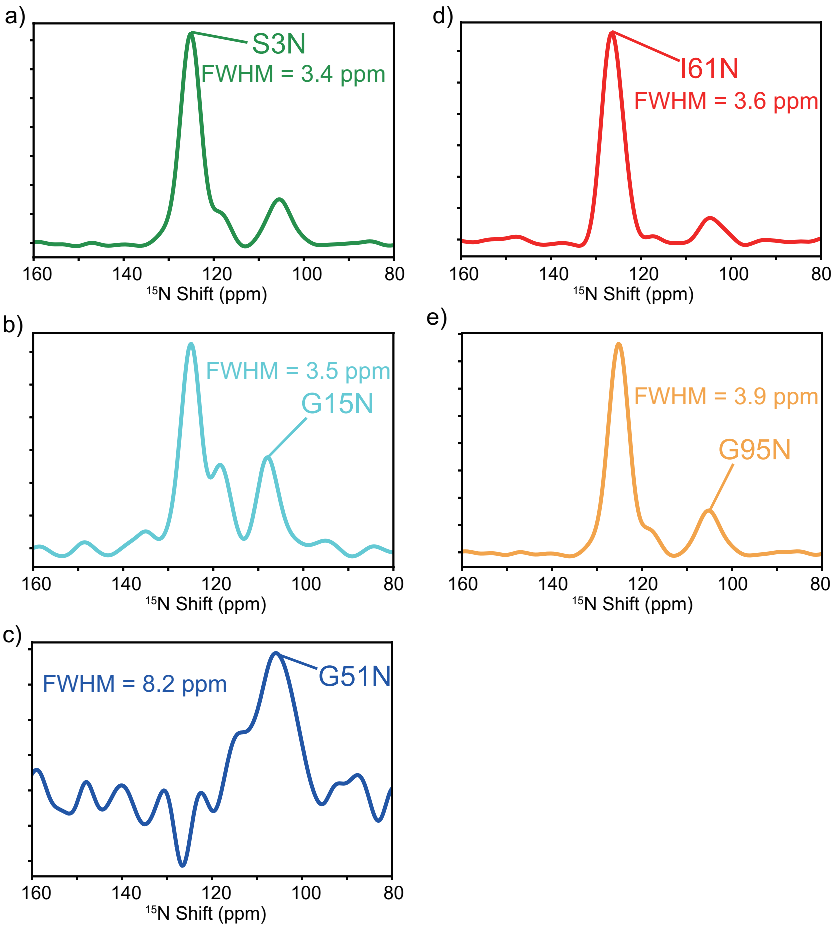


**Fig. S3.** The 1D slices of S3N (a), G15N (b), G51N(c), I61N(d) and G95N (e) extracted from 2D NC' DCP correlation spectrum of IG-DHFR (**Fig. 2b**) at the C chemical shifts of their corresponding peak centers show that the line broadening is site-specific.

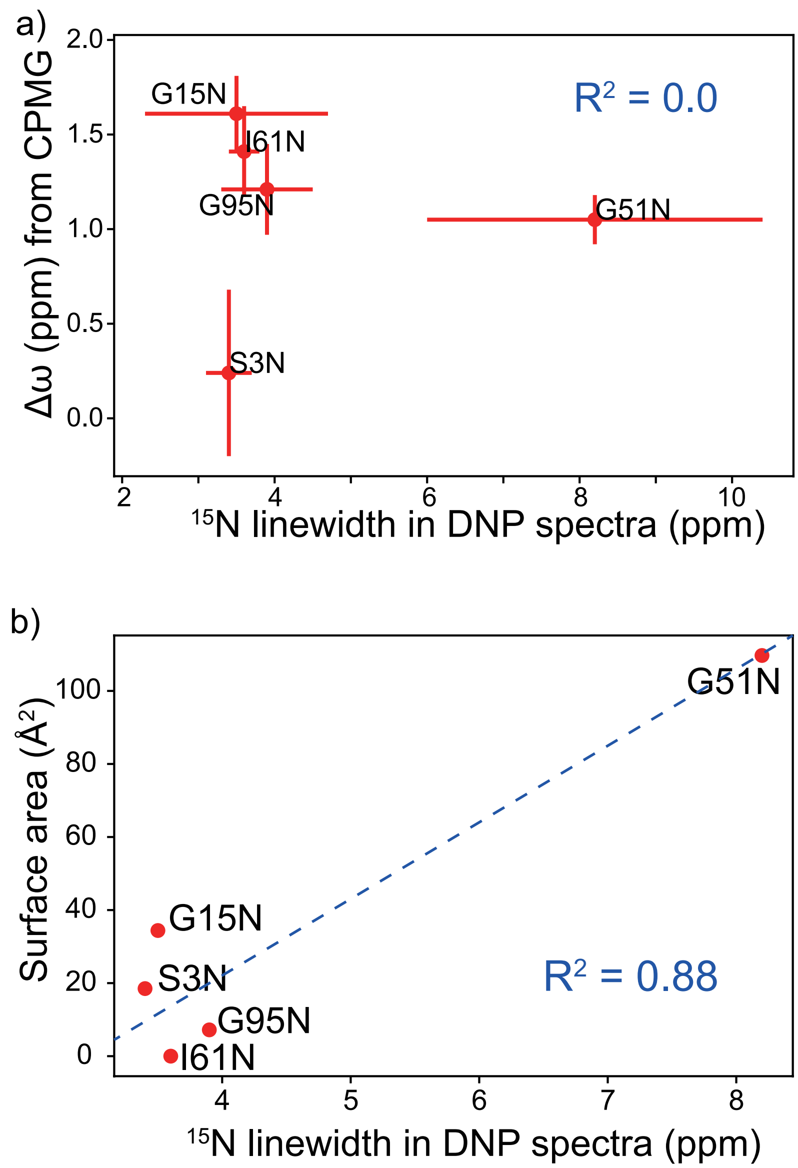


**Fig. S4.** (a) No correlation was observed between the N linewidth **(Fig. S2)** observed at low temperature and the chemical shift difference between ground state and active state at room temperature obtained from CPMG relaxation dispersion experiments at 300 K. The CPMG experiments were performed on uniformly enriched but otherwise similar samples. The N linewidth ranges from 3.4 ppm to 8.2 ppm while most of $\Delta\omega$ of each residue is within 2 ppm. (b) The correlation between the N linewidth **(Fig. S2)** observed at low temperature and the surface area at each residue in the DHFR:TMP X-ray crystallography structure (11) is shown. It is possible that there is some relationship; for example, Gly51 is more exposed to solvent compared to the other 4 residues and it exhibits the broadest linewidth. Although the overall coefficient of determination was 0.88 for the 5 residues, it is mainly driven by the distinct linewidth and solvent accessibility of Gly51. The correlation between solvent accessibility and low temperature linewidth is not strong when the $^{15}$N linewidth is within 4 ppm, possibly because of errors in linewidth estimate, peak overlaps and errors in solvent accessibility estimates, or other factors that contribute to line broadening at cryogenic temperature.

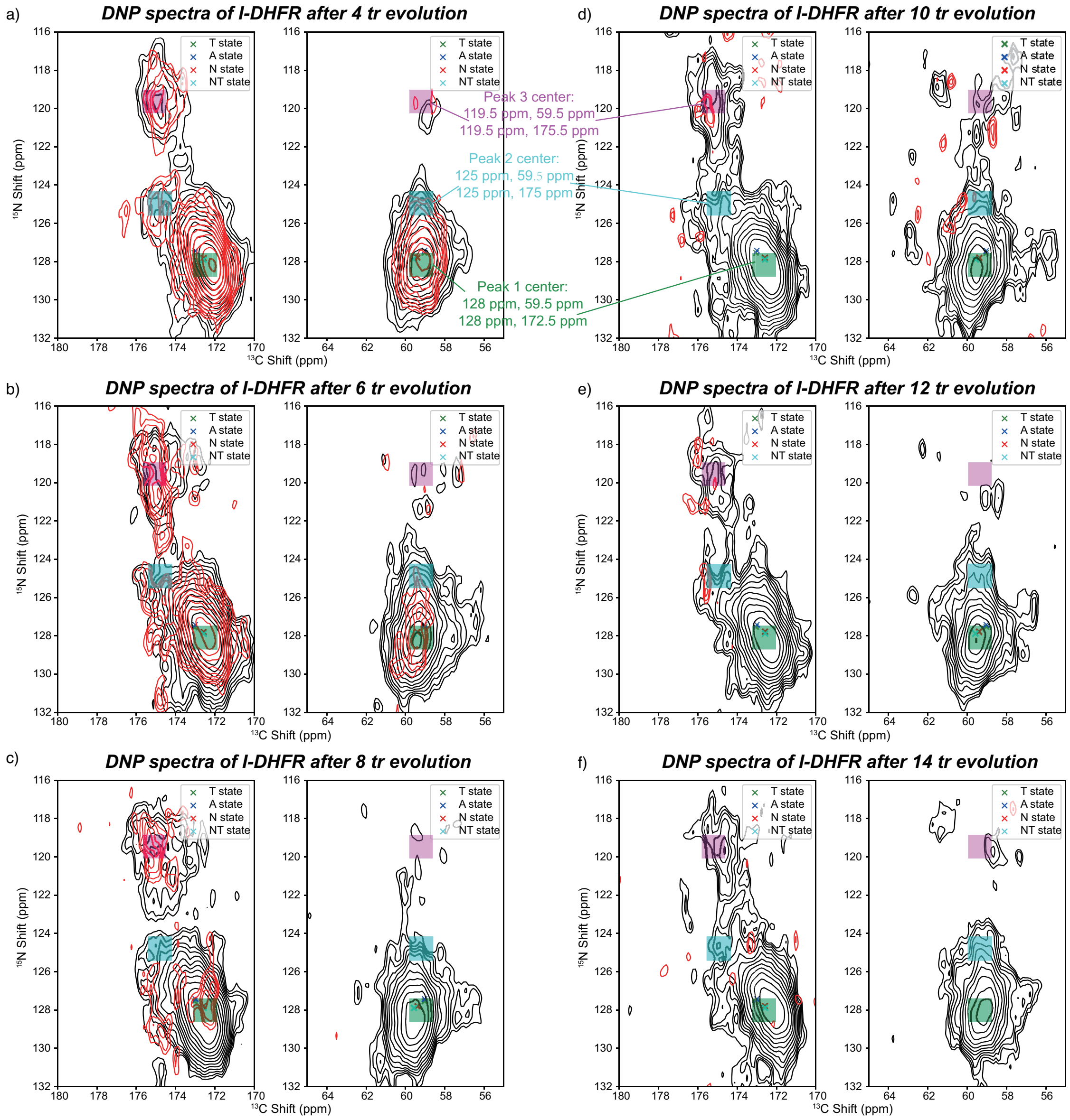


**Fig. S5.** 2D NCCN Ψ DQ dephasing $S_r$ (red) and $S_0$ (black) spectra at $4\tau r$ (a), $6\ \tau r$ (b), $8\ \tau r$ (c), $10\ \tau r$ (d), $12\ \tau r$ (e) and $14\ \tau r$ (f). The experiment details and spectra processing parameters are listed in **Table 2S**. The colored regions were determined as the Peak 1-3 as **Fig. 3b**, and the highest intensities within each peak region of C' and Cα are added to calculate the decay. Prior to taking $S_r/S_0$ ratio, the cross-peak intensities extracted from the $S_r$ and $S_0$ experiments were normalized per number of scans. The cross marker colored in green, blue, red and cyan represent the solution-state NMR chemical shifts of DHFR:TMP (T) state, Apo (A) state, DHFR:NADPH (N) state and DHFR:NADPH:TMP (NT) state at 300 K.

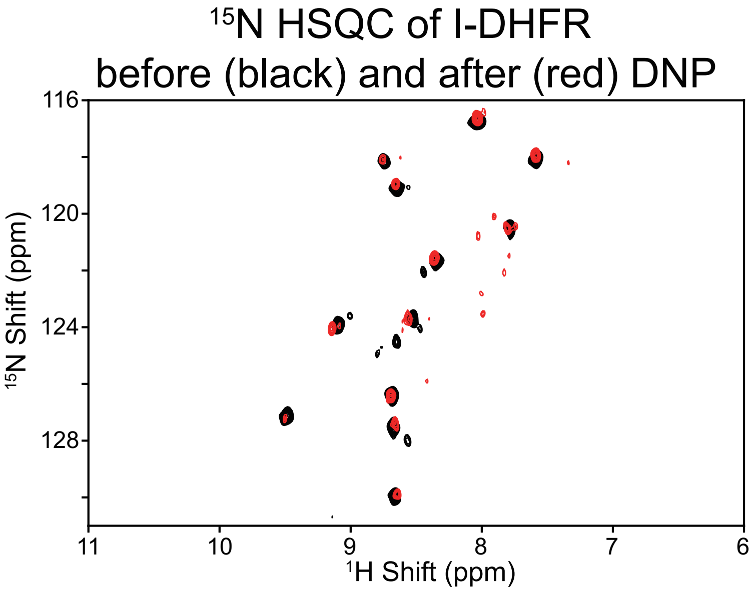


**Fig. S6.** $^{15}$N HSQC spectra of I-DHFR before (black) and after (red) the DNP experiments. All 12 Ile in the DHFR sequence were observed at the expected peak positions. The black spectrum was taken on a 700 MHz Bruker spectrometer. The red spectrum was taken on an 800 MHz Bruker spectrometer. Both spectra were taken at 300 K in HEPES buffer. The experimental details and processing parameters are listed in **Table 3S**. The protein concentration was 1 mM for the spectrum collected before DNP and 0.15 mM after DNP. The number of scans for black spectrum is 16 and for red is 128. The signal-to-noise at the peak of Ile61H-N is 33 before the DNP and 11 after that. Considering the protein concentration signal averaging and intrinsic sensitivity of the two instruments, the spectra have similar normalized signal-to-noise and 85% of protein was successfully recovered after the DNP experiments.

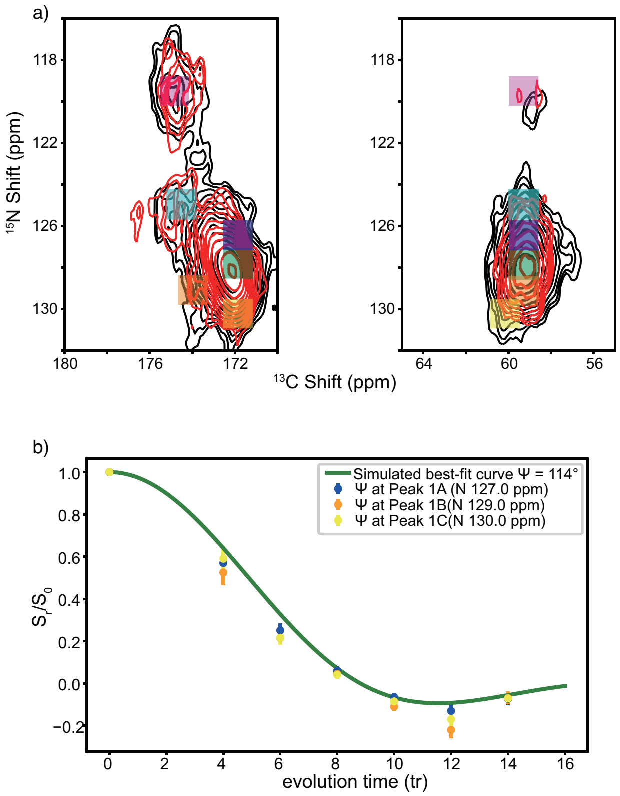


**Fig. S7.** Comparison of torsion angle measurements at three peak positions within Peak 1 centered at 128 ppm. (a) Three peaks, Peak 1A, Peak 1B and Peak 1C with different N and C peak positions (Peak 1A: N 127 ppm, C' 172 ppm, Cα 59.5 ppm; Peak 1B: N 129 ppm, C' 174 ppm, Cα 59.5 ppm; Peak 1C: N 130 ppm, C' 172 ppm, Cα 60 ppm) are indicated in blue, orange and yellow, respectively. (b) The intensity ratios of each peak were calculated similarly as was done for the three main peaks (**Fig. 4**) and their decay trends are very close.

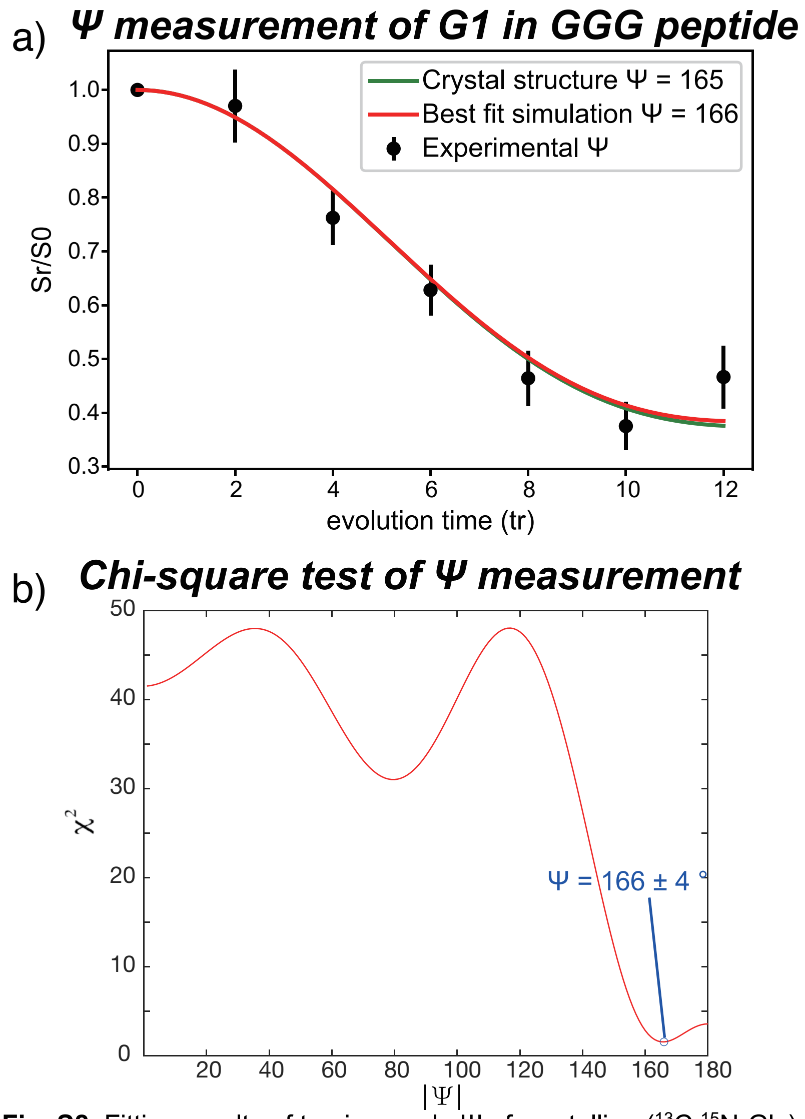


**Fig. S8.** Fitting results of torsion angle Ψ of crystalline ($^{13}C$,$^{15}N$-Gly)-($^{15}N$-Gly)-Gly·HCl. Scatter points in (a) are normalized C' and Cα intensities from $S_r$ and $S_0$ spectra collected using the NCCN Ψ DQ dephasing experiment (1, 12) (illustrated in **Fig. S1**). Red curve in (a) is the fitting result from Chi-square test, and green curve corresponds to the dephasing of the first G in GGG from X-ray crystallography, which is 165° (13). (b) Fitting result is the angle with the smallest Chi-Square value, which is 166°. The error range is determined by the 90% confidence level in the Chi-square fit (8).

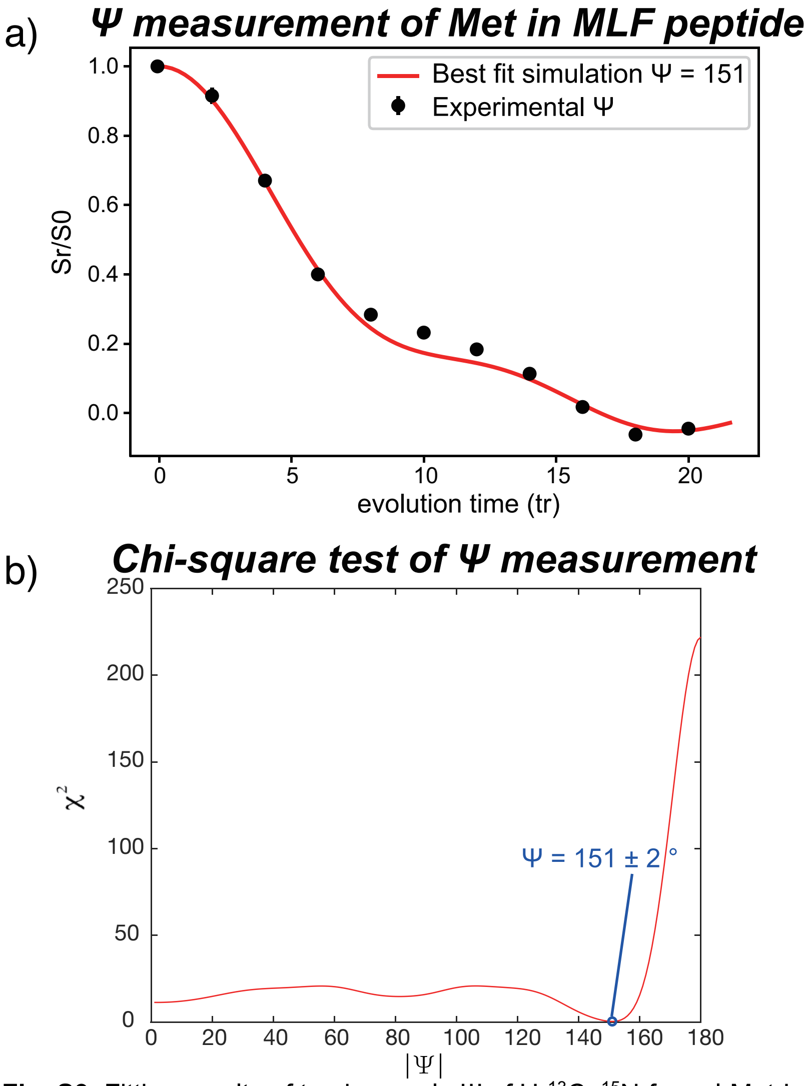


**Fig. S9.** Fitting results of torsion angle Ψ of U-$^{13}$C, $^{15}$N-formyl-Met-Leu-Phe-OH. Scatter points in (a) are normalized the average of C' and Cα intensities from $S_r$ and $S_0$ spectra collected using the NCCN Ψ DQ dephasing experiment (1, 12) (illustrated in **Fig. S1**). Red curve in (a) is the fitting result from Chi-square test, and it matches with the Ψ of Met in MLF from X-ray crystallography, which is 151°(14). (b) Fitting result is the angle with the smallest Chi-Square value, which is 151°. The error range is determined by the 90% confidence level in the Chi-square fit(8).

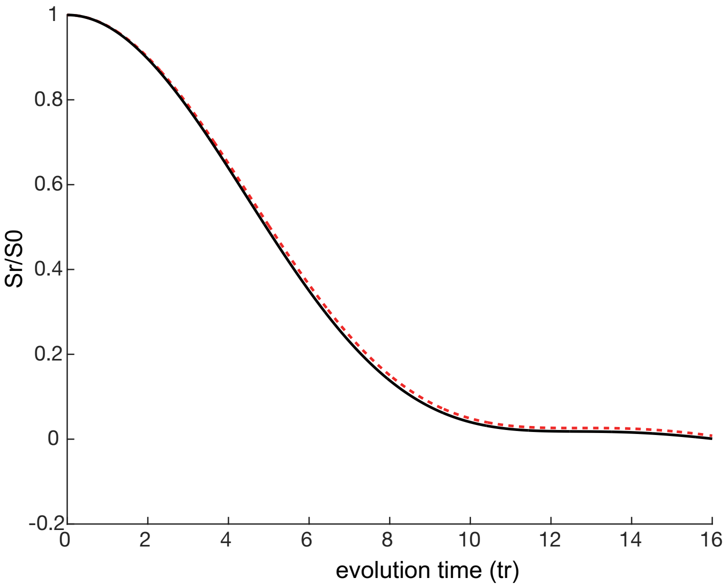


**Fig. S10.** Simulation results of Ψ = 140° with/without two-bond dipolar couplings effect. Both simulations were performed in MATLAB_R2019a using spinach_2.1.4400 (6). The result with 2 two-bond dipolars, which is $C'_i$-$N_i$ and $C\alpha_i$-$N_{i+1}$, is shown in black curve, while the red dash curve is the curve with only one-bond dipolar couplings into consideration. This result shows that the one-bond dipolar is the dominant contribution to this torison angle decay curve. This indicates that the two-bond dipolars have little effects on the dephasing.

**Table S1.** Protein residue flexibility analysis comparing the torsion angle variations among different crystals. Torsion angle variations in between DHFR:TMP state and all 6 conformations (15) in DHFR catalytic cycle are summarized. I14-G15 shows the largest mobility. I2-S3 and I60-I61 were categorized as rigid sites, while the rest three were mobile sites.

| Residue pair | Torsion angle | Secondary structure in DHFR:TMP X-ray structure | | DHFR: NADPH | DHFR: NADPH :DHF | Tran-sition state | DHFR: NADP+ : THF | DHFR: THF | DHFR: NADPH :THF | | Torsion angle variation range |
|---|---|---|---|---|---|---|---|---|---|---|---|
| | | | DHFR: TMP | DHFR: NADPH (1rx1) | DHFR: NADP+ :FOL (1ra2) | DHFR: NADPH :MTX (1rx3) | DHFR: NADP+ :ddTHF (1rc4) | DHFR:d dTHF (1rx5) | DHFR: NADPH :ddTHF (1rx6) | Apo (5dfr) | |
| I2-S3 | I2 psi | β-sheet | 131 | 133 | 134 | 136 | 133 | 134 | 135 | 133 | 5 |
| I14-G15 | I14 psi | coil | 1.7 | -13 | -11 | -20 | 157 | 150 | 161 | -26 | 181 |
| I50-G51 | I50 psi | α- helix | -44 | -32 | -42 | -26 | -38 | -37 | -41 | 143 | 18 |
| I60-I61 | I60 psi | β-sheet | 118 | 120 | 120 | 124 | 122 | 130 | 125 | 175 | 12 |
| I94-G95 | I94 psi | coil | 18 | 7.2 | 17 | 3.2 | 19 | 3.7 | 19 | -46 | 15.8 |

**Table S2.** Detailed acquisition and processing parameters for presented solid-state NMR spectra. All spectra listed here were acquired on a Bruker ACANCE spectroscopic. All the pulse sequence used are available in the Solids Pulse Sequence Library at New York Structural Biology Center. All the solid-state NMR experiments were performed at ~100 K with DNP. All spectra were processed using Topspin. The estimated noise was defined as the RMS of a non-peak region and was used for signal to noise calculation.

| Spectrum | N(Co)Cx | | NCo | | NCCN psi | |
|---|---|---|---|---|---|---|
| Pulse sequence | cpF3_dbcp_darr.kjf | | dbcp.kjf2 | | cp_dbcp_spc5_grif.xyi | |
| Figure(s) | 2 | | 3 | | 3, S5 | |
| Sample | IG-DHFR | | I-DHFR | | I-DHFR | |
| DNP enhancement | 90x | | 75x | | 75x | |
| MAS | 9 kHz | | 10 kHz | | 10 kHz | |
| DCP mixing time (p30) | 4 ms | | 4 ms | | 4 ms | |
| DCP efficiency | 50% | | 44% | | 44% | |
| DQ excitation / reconversion power (plw11) | - | | - | | 18.5 W (50 kHz) | |
| Dimension | F1 | F2 | F1 | F2 | F1 | F2 |
| Nucleus | C | N | C | N | C | N |
| Acq. time (ms) | 6 | 6 | 13 | 10 | 26 | 20 |
| π pulse length | - | - | - | - | 6.01 us | 8.06 us |

| | | | | | | |
|---|---|---|---|---|---|---|
| lowest contour (*RMS noise level) | 5 | | 5 | | 5 (Fig. 3); 4 (Fig. S5a-b); 3 (Fig. S5c-f) | |
| multiplicative factor for other levels | 1.3 | | 1.3 | | 1.3 (Fig. 3); 1.2 (Fig. S5) | |
| LB (type) | GM | GM | GM | GM | EM | EM |
| LB (Hz) | 150 | 150 | 50 | 50 | 50 | 50 |

**Table S3.** Detailed acquisition and processing parameters for presented solution-state NMR spectra. All spectra listed here were acquired on a Bruker ACANCE spectroscopic. All the pulse sequence used in solution-state NMR experiments are from standard Bruker library.

| Spectrum | N-HSQC | | N-HSQC | | H(N)C' | | H(NC')Ca | |
|---|---|---|---|---|---|---|---|---|
| Pulse sequence | AR_hsqcetfpf3gpsi2 | | AR_hsqcetfpf3gpsi2 | | AR_hncogp3d | | AR_hncocagp3d | |
| Spectrometer | 700 MHz | | 800 MHz | | 700 MHz | | 700 MHz | |
| Figure(s) | S2 | | S6 | | S2 | | S2 | |
| Sample | I-DHFR | | recovered I-DHFR | | I-DHFR | | I-DHFR | |
| Scans | 16 | | 128 | | 256 | | 256 | |
| FIDs | 128 | | 128 | | 64 | | 64 | |
| Dimension | F1 | F2 | F1 | F2 | F1 | F2 | F1 | F2 |
| Nucleus | H | N | H | N | H | C' | H | Ca |
| Window Functions | QSINE | QSINE | QSINE | QSINE | QSINE | QSINE | QSINE | QSINE |
| SSB | 2.5 | 2.5 | 2.5 | 2.5 | 2 | 2 | 2 | 2 |
| Signal to Noise | 33 | | 11 | | 6 | | 11 | |

**Table S4.** Average peak positions of Ile61N, Ile60C’ and Ile60Cα weighted by peak intensities at Peak1, Peak2, Peak3. The comparison between the solution-state NMR chemical shifts at 300 K of Ile61N, Ile60C’ and Ile60Cα and the averaged peak positions at 100 K shows a good match.

| Peaks | Peak 1 | Peak 2 | Peak 3 | Average over peak intensities | **Average over peak volumes** | **X-ray structure** | **Solution-state chemical shifts (ppm)** |
|---|---|---|---|---|---|---|---|
| Peak intensity at 100 K | 3.9E+05 | 1.0E+05 | 7.8E+04 | - | **-** | **-** | **-** |
| Peak volume at 100 K | 2.0E+07 | 3.9E+06 | 6.5E+06 | - | **-** | **-** | **-** |
| Psi angle (degrees) | 114.0 | 150.0 | 164.0 | 127.3 | **129.5** | **118.0** | **-** |
| N Peak position (ppm) | 128.0 | 125.0 | 119.5 | 126.3 | **125.8** | **-** | **127.5** |
| C' Peak position (ppm) | 172.0 | 175.0 | 175.5 | 173.0 | **173.1** | **-** | **173.0** |
| CA Peak position (ppm) | 59.5 | 59.5 | 59.5 | 59.5 | **59.5** | **-** | **59.0** |

**SI References:**


1. V. Ladizhansky, C. P. Jaroniec, A. Diehl, H. Oschkinat, R. G. Griffin, Measurement of multiple ψ torsion angles in uniformly 13C,15N-labeled α-spectrin SH3 domain using 3D 15N-13C--13C-15N MAS dipolar-chemical shift correlation spectroscopy. *J. Am. Chem. Soc.* **125**, 6827–6833 (2003).
2. M. Baldus, A. T. Petkova, J. Herzfeld, R. G. Griffin, Cross polarization in the tilted frame: Assignment and spectral simplification in heteronuclear spin systems. *Mol. Phys.* **95**, 1197–1207 (1998).
3. J. Schaefer, R. . McKay, E. O. Stejskal, Double-cross-polarization NMR of solids. *J. Magn. Reson.* **34**, 443–447 (1979).
4. M. Hohwy, C. M. Rienstra, R. G. Griffin, Band-selective homonuclear dipolar recoupling in rotating solids. *J. Chem. Phys.* **117**, 4973–4987 (2002).
5. T. Gullion, J. Schaefer, Rotational -echo double-resonance NMR. *J. Magn. Reson.* **81**, 196–200 (1989).
6. H. J. Hogben, M. Krzystyniak, G. T. P. Charnock, P. J. Hore, I. Kuprov, Spinach - A software library for simulation of spin dynamics in large spin systems. *J. Magn. Reson.* **208**, 179–194 (2011).
7. W. Modell, *CRC Handbook of Tables for Probability and Statistics* (1966).
8. C. Garland, J. Nibler, D. Shoemaker, Experiment 25 Surface Tension of Solutions. *Exp. Phys. Chem.*, 299–308 (2009).
9. P. R. Costa, J. D. Gross, M. Hong, R. G. Griffin, Solid-state NMR measurement of Ψ in peptides: A NCCN 2Q-heteronuclear local field experiment. *Chem. Phys. Lett.* **280**, 95–103 (1997).
10. X. Feng, *et al.*, Direct determination of a molecular torsional angle by solid-state NMR. *Chem. Phys. Lett.* **257**, 314–320 (1996).
11. R. Rogawski, Dynamic Nuclear Polarization with Biradical Affinity Reagents (2018).
12. P. R. Costa, J. D. Gross, M. Hong, R. G. Griffin, Solid-state NMR measurement of Ψ in peptides: A NCCN 2Q-heteronuclear local field experiment. *Chem. Phys. Lett.* **280**, 95–103 (1997).
13. V. L. and E.Subramanian, Glycyl-glycyl-glycine hydrochloride, C6H11N3O4.cnt. 11,561. (1982).
14. C. M. Rienstra, *et al.*, De novo determination of peptide structure with solid-state magic-angle spinning NMR spectroscopy. *Proc. Natl. Acad. Sci.* **99**, 10260–10265 (2002).
15. M. R. Sawaya, J. Kraut, Loop and subdomain movements in the mechanism of Escherichia coli dihydrofolate reductase: Crystallographic evidence. *Biochemistry* **36**, 586–603 (1997).
